# Gb3 trisaccharide-bearing exosomes as a novel neutralizer for Shiga toxin type 1

**DOI:** 10.1101/2024.10.26.620411

**Authors:** Krzysztof Mikołajczyk

## Abstract

Shiga toxin types 1 (Stx1) and 2 (Stx2), produced by Shiga toxin-producing *Escherichia coli* (STEC) and *Shigella dysenteriae*, are key virulence factors responsible for severe foodborne diseases, such as hemorrhagic colitis and hemolytic uremic syndrome (HUS). The receptors for Stxs are Gb3 and P1 glycotope, which contain the Galα1→4Gal epitope and are synthesized by human α1,4-galactosyltransferase (A4galt). Stx-related infections pose a global public health challenge, owing to the limited therapeutic options due to the restricted use of antibiotics. Therefore, there is an urgent need to develop novel therapeutic strategies. This study proposes an innovative strategy utilizing exosomes derived from CHO-Lec2 cells, which were modified with Functional-Spacer-Lipid (FSL) conjugates bearing the Gb3 carbohydrate epitope (exo-Gb3-FSL). Flow cytometry analysis confirmed the presence of Galα1→4Gal disaccharides on exo-Gb3-FSL constructs, enabling them to bind Stx1. Moreover, using CHO-Lec2 cells evaluated the ability of exo-Gb3-FSL agents to bind Stx1 and protect these cells from Stx1-mediated cytotoxicity. For Stx1-treated CHO-Lec2 cells, increased cell survival was observed when using 25 µM exo-Gb3-FSL constructs, compared to control cells. These findings highlight the potential of exosome-based anti-Stx1 agents as promising alternatives to conventional therapies. This innovative strategy may provide novel directions for studies on Stx1 neutralization, offering a valuable strategy for the treatment of Stx-related diseases.

**Highlights:**

- Exo-Gb3-FSL constructs hold Stx1 binding Galα1→4Gal epitope.
- Stx1 was effectively bound by exo-Gb3-FSL constructs.
- Exo-Gb3-FSL protected CHO-Lec2 cells from Stx1.
- Exo-Gb3-FSL is a promising anti-Stx1 agent.

## 1 Introduction

Shiga toxins (Stxs) are produced by foodborne pathogens *Shigella dysenteriae* serotype 1, enterohaemorrhagic *Escherichia coli* (EHEC) strains and Stx-producing *E. coli* (STEC). STEC produces two Stx types: Stx1 is 98 % identical to the Shiga toxin produced by *Shigella dysenteriae*, whereas Stx2 shares approximately 65 % similarity in amino acid composition, but exhibits greater heterogeneity between subtypes [1, 2, 3]. Stxs display an AB_5_ structure, comprising one catalytic A domain and a pentamer of B subunits interacting with the Galα1→4Gal epitope, found in the Gb3 glycosphingolipid (GSL). Gb3 (Galα1→4Galβ1→4Glc-Cer, also called globotriaosylceramide, P^k^) is a glycosphingolipid antigen belonging to the human P1PK blood group system [4]. Recent studies have identified a glycoprotein (GP)-based receptor, known as the P1 glycotope, which is capable of Stx1 binding [5, 6]. Both Gb3 and P1 glycotope are synthesized by human α1,4-galactosyltransferase (A4galt) and have been extensively studied using Chinese hamster ovary (CHO-Lec2) cells to elucidate their interactions with Stxs [5, 7].

STEC infections are associated with severe gastrointestinal and kidney-related aberrations, including hemorrhagic colitis and hemolytic uremic syndrome (HUS). HUS is a severe and potentially life-threatening complication characterized by thrombocytopenia, hemolytic anemia, and acute renal failure, and remains a major cause of kidney damage and neurological complications in humans. STEC causes >20,000 infections annually, with clinical manifestations ranging from bloody diarrhea and hemorrhagic colitis to hemolytic uremic syndrome (HUS) [8, 9]. Current therapeutic approaches for STEC and Stx-related diseases are limited because antibiotic treatment can exacerbate the condition by inducing bacterial lysis and increasing toxin release. This effect highlights the urgent need for novel therapeutic strategies to avoid antibiotic usage. No specific therapeutics are recommended for treating Stxs-related diseases, allowing mainly supportive care of the patients [9]. In light of these challenges in using conventional antimicrobial treatments for Stx-related diseases, recent efforts have been directed towards developing novel approaches, including Stx neutralizers and inhibitors (reviewed in [9]).

One of the promising therapeutic alternatives against Stx-related diseases are Stx neutralizers [9]. In this study, it is proposed the use of the FSL conjugate as a carrier of the Stx-binding Galα1→4Gal disaccharide. FSL conjugates have an amphipathic chemical structure that allows their spontaneous and stable incorporation into membrane bilayers of various entities, including cells, liposomes, and extracellular vesicles, such as exosomes. Once incorporated, FSL conjugates can present specific glycan epitopes on the surface of these membranes. Structurally, FSL consists of three components: a functional head group, spacer, and lipid tail [10, 11]. Notably, previous studies have demonstrated the efficacy of Gb3-containing FSL conjugates in preventing viral infections, such as HIV [12].

Here, the synthetic analog of Gb3 containing a Galα1→4Galβ1→4Glcβ-R trisaccharide epitope conjugated via an O(CH2)3NH spacer (SA1) to an activated adipate derivative of dioleoylphosphatidylethanolamine (DOPE) (L1) served as the anti-Stx1 agent (Fig. 1A). This innovative approach involved incorporating the Gb3-FSL conjugate into exosomes isolated from CHO-Lec2 cells to create exo-Gb3-FSL constructs capable of binding Stx1. Exosomes are small extracellular vesicles with a phospholipid bilayer [13] that provide an ideal platform for embedding the Gb3-FSL conjugate into their membranes. This creates exosome particles adorned with Gb3-mimicking molecules on their surface. These exo-Gb3-FSL constructs were then analyzed for their potential anti-Stx1 properties using A4galt-expressing CHO-Lec2 cells (Stx1-sensitive), allowing for the assessment of their ability to neutralize Stx1 binding.

**Fig. 1.**
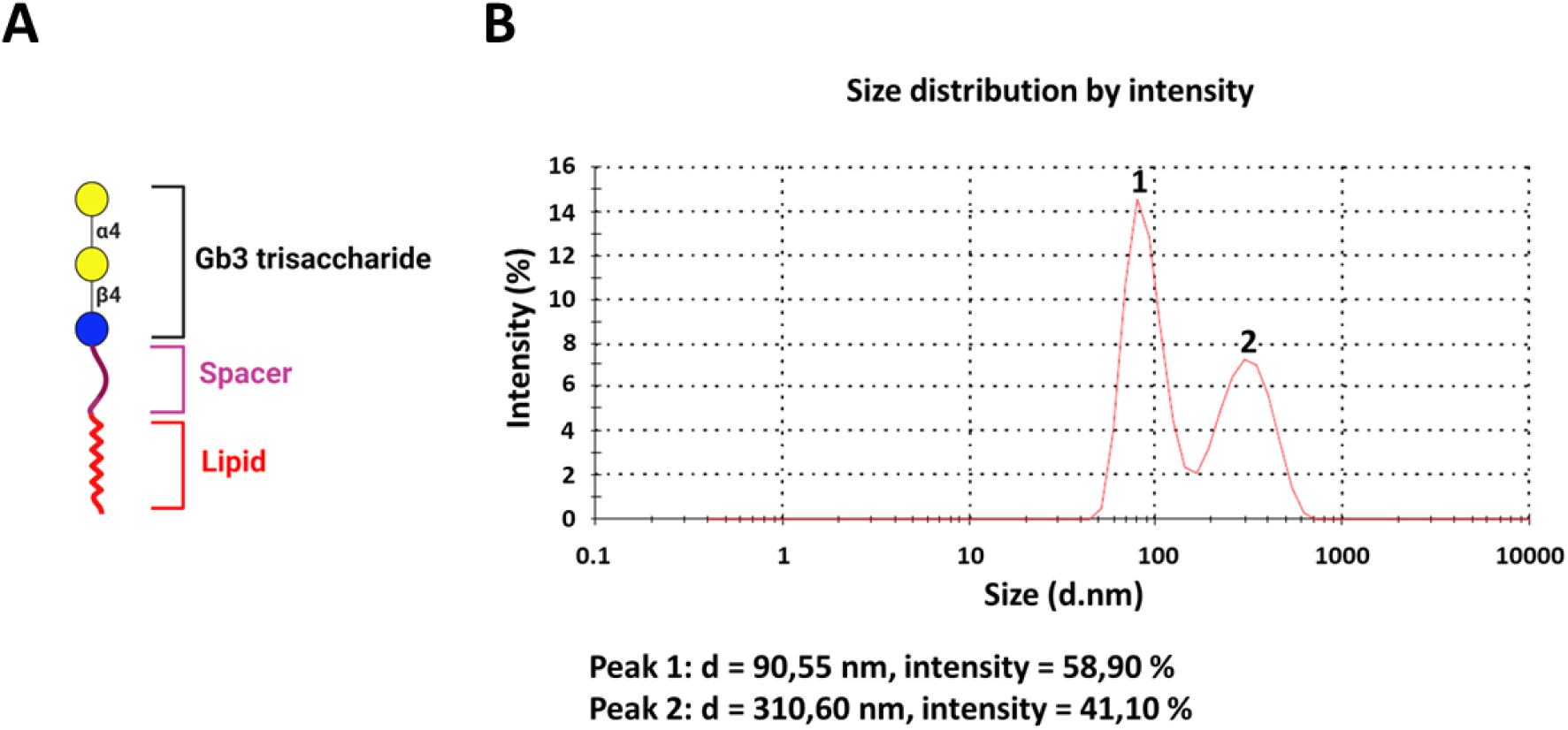
Exosomes isolated from CHO-Lec2 cells were incubated with Gb3 trisaccharide-containing (Galα1→4Galβ1→4Glcβ-R) FSL conjugate (designated as exo-Gb3-FSL). Exo-Gb3-FSL consists of (1) Gb3 trisaccharide, (2) O(CH_2_)_3_NH spacer and (3) activated adipate derivative of dioleoylphosphatidylethanolamine lipid (**A**). The particle size distribution of CHO-Lec2 NAT-derived exosomes was evaluated using the DLS Zetasizer (Malvern) (**B**).

## 2 Material and methods

### 1.1 Cell culture

CHO-Lec2 cells were obtained from the American Type Culture Collection (Rockville, MD, USA). The cells were grown and maintained in a humidified incubator with 5% CO2 at 37°C using DMEM/F12 medium (Thermo Fisher Scientific, Inc., Waltham, MA, USA) with 10% fetal bovine serum (Thermo Fisher Scientific, Inc., Waltham, MA, USA) and Pen-Strep (Thermo Fisher Scientific, Inc., Waltham, MA, USA). CHO-Lec2 cells transfected with the mutein variant of human α1,4-galactosyltransferase were obtained as described previously [5].

### 1.2 Exosome isolation and characterization

Untransfected CHO-Lec2 cells (NAT) were plated in cell culture flasks containing completed DMEM/F12 medium (Thermo Fisher Scientific, Inc., Waltham, MA, USA). The next day, the medium was replaced with FBS-depleted DMEM/F12 medium after washing three times with phosphate-buffered saline (PBS). After a 48-h incubation period, the medium was harvested for exosome isolation, when cells reached approximately 90% confluence. The collected medium was sequential centrifugated at 600 × *g* for 10 min, 2000 × *g* for 15 min, and 10,000 × *g* for 30 min to remove cellular debris and larger particles. The remaining supernatant was then subjected to two rounds of ultracentrifugation at 100,000 × *g* for 70 min at 4 °C to isolate exosomes, following the protocol [14]. The exosome pellet was resuspended in 100 μL of PBS and stored in Protein LoBind tubes (Eppendorf, Hamburg, Germany) at either 4 °C or −80 °C for further analysis. Dynamic light scattering (DLS) (Malvern Zetasizer, Malvern Instruments, UK) was used to assess the size distribution and purity of the isolated exosomes, with samples diluted 1:1 in PBS. All steps were performed at 4 °C to preserve the exosome integrity.

### 1.3 Flow cytometry

CHO-Lec2 cells, both untransfected (NAT) and transfected with a mutein variant of human A4galt (A4G) were trypsinized, washed, and subsequently incubated with anti-P1 (clone 650) antibody and Stx1B-AF488 as described in [5]. The exosomes-Gb3-FSL constructs were bound to aldehyde/latex beads, 4% w/v, 4 μm (Thermo Fisher Scientific, Inc., Waltham, MA, USA), and blocked with 100 mM glycine solution (Sigma-Aldrich, St. Louis, MO) and incubated with anti-P1 (clone 650), anti-CD81 (clone 2F2) antibodies as well as with Stx1B-AF488 (conjugated with AlexaFluor 488 with Alexa Fluor™ 488 Protein Labeling Kit, Thermo Fisher Scientific, Inc., Waltham, MA, USA). To evaluate Stx1B-AF488 sequestration, the exo-Gb3-FSL constructs were incubated with Stx1B-AF488 overnight at 4 °C, then washed three times and added to CHO-Lec2 NAT and A4G cells. The conjugated beads and CHO-Lec2 cells were analyzed by flow cytometry in the proper cytometer settings (FACSCalibur, BD Biosciences, Franklin Lakes, NJ, USA).

### 1.4 Cell viability assay

The exosome-containing Gb3-FSL conjugates were prepared by coincubation of exosomes (100 µl) isolated from untransfected CHO-Lec2 and Gb3-FSL conjugates (kindly provided by Nicolai Bovin, Kode Biotech Ltd., Auckland, New Zealand) at 37 °C by 1 h with agitation (350 rpm). To remove unbound Gb3-FSL conjugates, the samples were ultracentrifuged at 100,000 × *g* for 70 min, resuspended in PBS and used for cytotoxicity assay.

The cell viability was evaluated using the CellTiter 96 AQueous One Solution Assay (Promega, Madison, WI) as described in [5]. Briefly, CHO-Lec2 NAT and CHO-Lec2 A4G cells were seeded (2 × 10^4^ cells/mL) in 96-well plates (Wuxi NEST Biotechnology Co, Ltd, China) in a complete DMEM/F12 medium (Thermo Fisher Scientific, Inc., Waltham, MA, USA). After 24 h the medium was replaced with serum-free DMEM/F12 containing varying concentrations of Stx1 holotoxin (0, 0.2, 0.4, 0.6, 0.8 and 1.0 ng/ml) (Sigma-Aldrich, St. Louis, MO) and exo-Gb3-FSL constructs (0, 5, 10, 15, 20, 25, 50, 75 and 100 µM). After treatment with toxins, the morphology of the cells will be assessed and MTS tetrazolium compound (CellTiter 96 AQueous One Solution Assay, Promega, Madison, WI) was added. The plates were incubated in a humidified, CO_2_ atmosphere for 4 h, and the absorbance at 490 nm was recorded using an ELISA plate reader (PerkinElmer 2300 EnSpire Multilabel Reader). The background absorbance registered at zero cells/well was subtracted from the data. The absorbance of wells incubated in medium without Stx was taken as 100% of cell viability. Each experiment was repeated at least three times.

## 3. Results

### 3.1 Characterization of CHO-Lec2-derived exosomes

For exosome isolation, untransfected CHO-Lec2 cells (NAT) were selected, based on their lack of endogenous expression of A4galt and deficiency of the CMP-sialic acid transporter. These features ensure that CHO-Lec2 is incapable of sialylation, which may compete with α-galactosylation during N-glycans synthesis [15, 16]. Additionally, extracellular vesicles, including exosomes, produced by CHO-Lec2 NAT cells are predicted to not contain glycolipids and glycoproteins terminated with the Galα1→4Gal epitope, which binds Stxs. Thus, CHO-Lec2 NAT cells seem to be a good cell model for producing exosomes deprived of Stx-binding receptors and for testing synthetic Gb3 receptor analogs, such as Gb3-FSL, to evaluate potential therapeutic capabilities.

Exosomes were isolated using a differential centrifugation and ultracentrifugation protocol as described previously [14]. To assess the size distribution of the isolated vesicles, dynamic light scattering (DLS) measurements were performed. The exosomes derived from CHO-Lec2 NAT cells exhibited an average diameter of approximately 162.1 nm, with two distinct subpopulations observed: one composed of smaller vesicles (∼90 nm) and another of larger vesicles (∼310 nm) (Fig. 1B). The exosome sample showed high homogeneity, characterized by the polydispersity index (PDI) of approximately 0.296.

For further characterization, CHO-Lec2 NAT-derived exosomes were immobilized onto 4 µm aldehyde/sulfate latex beads for flow cytometry analysis. It was found that latex microspheres-bound exosomes displayed surface expression of CD81, a well-known tetraspanin-related exosome marker (Fig. 2) [14]. These data confirmed the successful isolation of a homogeneous population of CHO-Lec2 NAT-derived exosomes, ensuring their suitability for subsequent investigations.

**Fig. 2.**
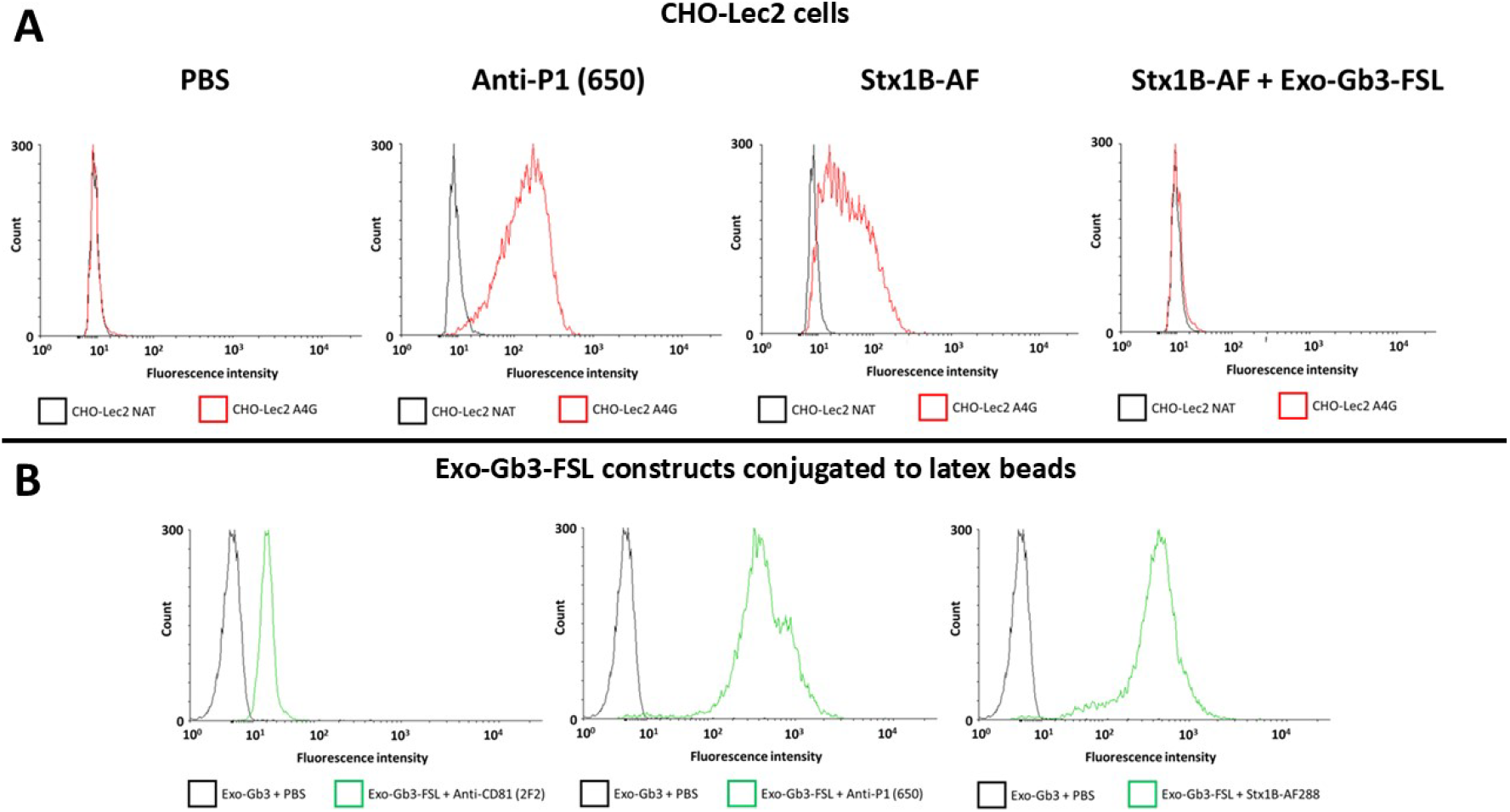
Flow cytometry analysis of (A) CHO-Lec2 cells transfected with vector encoding mutein variant (with p.Q211E substitution) of human α1,4-galactosyltransferase (A4G) or untransfected control (NAT) and (B) 4 µm aldehyde/sulfate latex beads captured CHO-Lec2 NAT-derived exosomes treated with Gb3-FSL conjugate. The binding of the anti-P1 (clone 650) antibody and Stx1B-AF488 (coincubated with exo-Gb3-FSL or not) was examined (**A**). Aldehyde/sulfate latex beads with captured exosomes treated or not with Gb3-FSL conjugate were evaluated using anti-CD81 (clone 2F2), anti-P1 (clone 650) and Stx1B-AF488 (**B**).

### 3.2 CHO-Lec2-derived exosomes can bind Stx1

To examine Stx1B binding, exosomes isolated from CHO-Lec2 NAT cells were modified with a Gb3-FSL conjugate, which carries a Gb3 trisaccharide linked to a diacyl lipid moiety capable of membrane bilayer incorporation. The obtained exosomes decorated with terminal Galα1→4Gal disaccharide (exo-Gb3-FSL) can interact with Stxs. To verify the presence of this epitope on the exosome surface and assess its ability to bind Stx1B-AF488 (fluorescently labeled Stx1B subunit), the exo-Gb3-FSL constructs were immobilized onto 4 µM aldehyde/sulfate latex beads and analyzed by flow cytometry. The binding of the anti-P1 antibody (clone 650), which binds Gb3 and P1 antigens, and Stx1B-AF488 to exo-Gb3-FSL-coated beads was confirmed (Fig. 2). Furthermore, coincubation of Stx1B-AF488 with exo-Gb3-FSL constructs coated on aldehyde/sulfate latex microspheres markedly reduced cell surface binding of Stx1B-AF488 to CHO-Lec2 A4G cells (Fig. 2). These results pinpoint the ability of exo-Gb3-FSL constructs to effectively sequester Stx1B.

### 3.3 Exo-Gb3-FSL constructs protects CHO-Lec2 A4G cells from Stx1 holotoxin

A cytotoxicity assay was performed to evaluate the efficacy of the exo-Gb3-FSL constructs as potential anti-Stx1 agents. In this assay, CHO-Lec2 NAT cells, which lack the Stx1 receptor, were not sensitive to Stx1 holotoxin exposure, whereas CHO-Lec2 A4G cells, which express Gb3 (a Stx1 receptor) were susceptible to Stx1-induced cytotoxicity. Exo-Gb3-FSL constructs, at concentrations ranging from 5 to 20 µM, provided a dose-dependent protective effect against Stx1 across various toxin concentrations. The cell viability of CHO-Lec2 A4G cells in the presence of 1 ng/ml Stx1 holotoxin ranged from 37% to 46% when co-incubated with these constructs (compared to 27% viability of the cells untreated with exo-Gb3-FSL constructs) (Fig. 3). In the presence of 25 µM exo-Gb3-FSL, CHO-Lec2 A4G cells exhibited approximately 56% viability, compared to 27% detected in untreated controls exposed to 1 ng/ml Stx1 (Fig 3). However, increasing the exo-Gb3-FSL concentration beyond 25 µM did not further enhance protection, but led to a slight reduction in cell viability to approximately 45% (also tested in the presence of 1 ng/ml Stx1) (Fig. 3). These results indicated that exo-Gb3-FSL constructs can serve as a Stx1 neutralizer, preventing the CHO-Lec2 A4G cells from Stx1-mediated cytotoxicity.

**Fig. 3.**
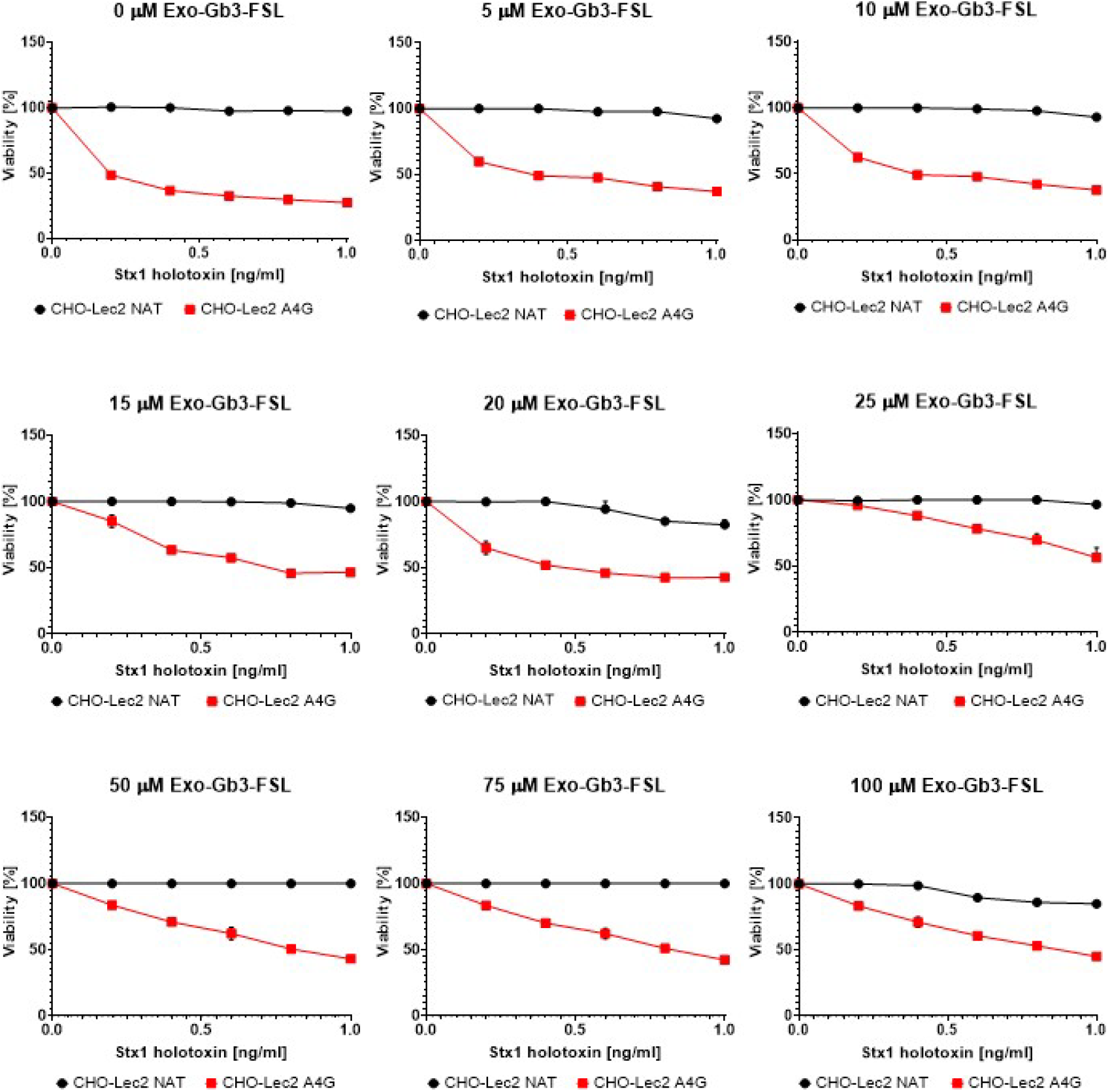
Shiga toxin type 1 cytotoxicity assay. The cell viability of CHO-Lec2 cells transfected with a vector encoding mutein variant (with p.Q211E substitution) of human α1,4-galactosyltransferase (A4G) and untransfected control (NAT) was evaluated. CHO-Lec2 cells were cultured in the presence or absence of exosomes containing various concentrations of the Gb3-FSL conjugate in a medium containing Stx1 holotoxin.

## 4 Discussion

In this study, it was demonstrated that exo-Gb3-FSL constructs serve as effective neutralizers of Stx1. Exosomes derived from CHO-Lec2 NAT cells, decorated with Gb3-FSL, mimicked the natural Gb3 trisaccharide, thereby preventing Stx1 from exerting cytotoxic effects on CHO-Lec2 A4G cells. To date, several Stx neutralizers have been developed, primarily focusing on inhibiting the internalization of Stx into target cells [9]. Among them, STARFISH [17, 18], Gb3 polymers (consisting of linear polymers of acrylamide with clustered Gb3 trisaccharides) [19], and Synsorb-Pk [20, 21, 22] were designed for local Stxs sequestration in the intestine, whereas SUPER TWIG is the only known agent capable of neutralizing circulating Stxs [23]. However, none of these neutralizers have demonstrated satisfactory efficacy and safety. For instance, Synsorb-Pk failed to show significant therapeutic effects compared with controls, with increased patient mortality reported [20, 21, 22]. Similarly, STARFISH, even when tested at a high dose of 25 µg/g in mice, showed no therapeutic effects, likely because of its short circulation half-life (less than 30 min) [17, 18]. Only, the modified STARFISH version showed promising *in vitro* and *in vivo* effectiveness against Stx1 and Stx2 [18].

Most of Stx neutralizers employ Gb3 trisaccharide analogs linked to various potential immunogenic matrices, such as dendrimers or acrylamide polymers [9]. Recently, a novel approach using Gb3-mExo, bovine milk-derived exosomes conjugated with stearic acid-based Gb3 derivatives, demonstrated effective Stx2 neutralization [24]. In addition, Duan et al. (2024) developed oligonucleotide-based aptamers that blocked Stx2 binding to Gb3 receptors in the cells [25]. The exo-Gb3-FSL constructs presented in this study represent a potentially efficient and biocompatible alternative to the previously evaluated Stx neutralizers. These constructs exploit the natural membrane integration of Gb3-FSL into exosomes (which can be isolated directly from patients in the future) without the potential side effects associated with using synthetic matrices [24]. The proposed protective effects of exo-Gb3-FSL may be a result of competition between Gb3 trisaccharide on exosomes and natural Gb3/P1 glycotope receptors expressed by the cells (Fig. 4). However, the precise molecular mechanisms underlying these interactions remain unclear and require further investigation.

**Fig. 4.**
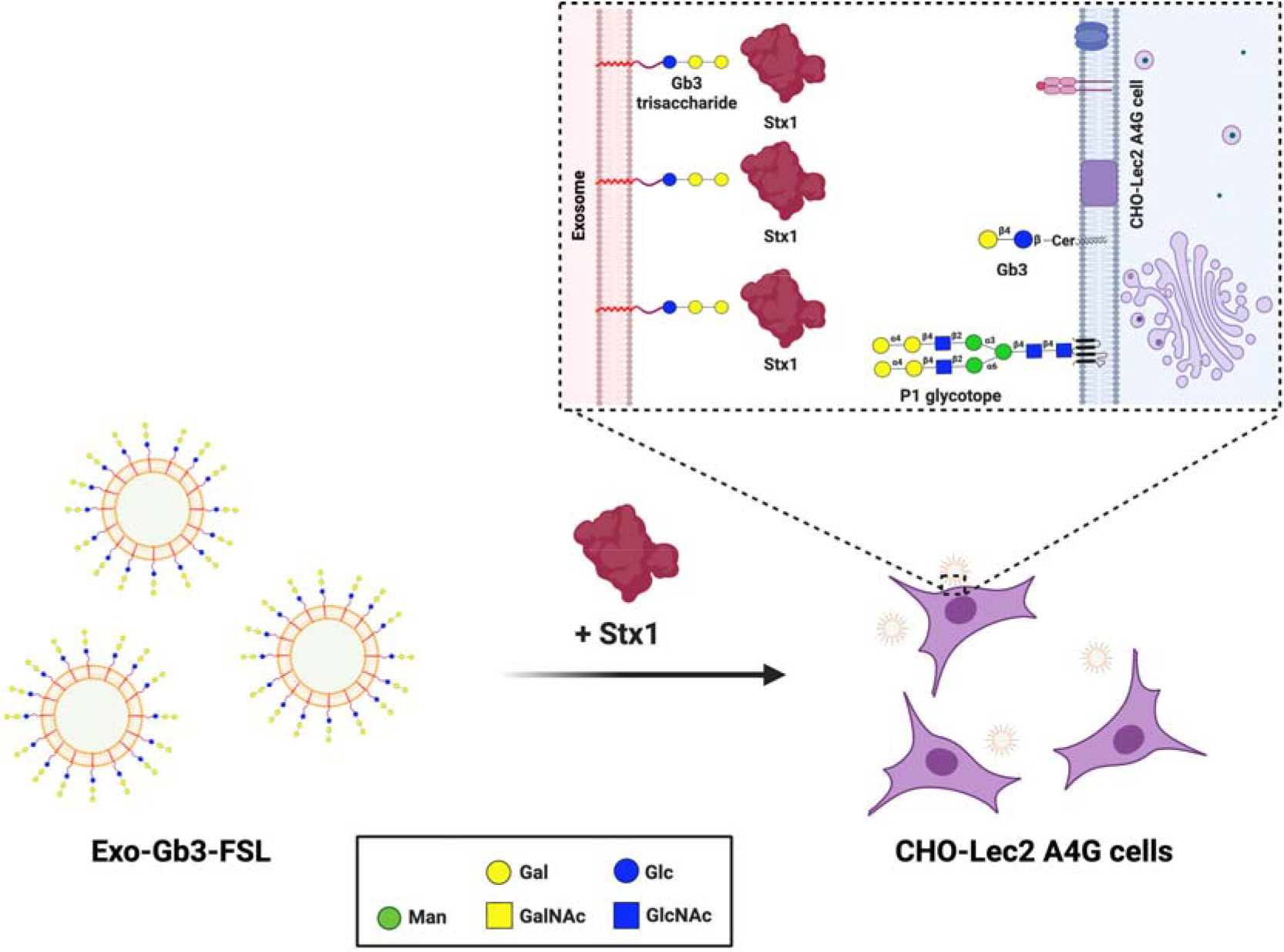
The proposed mechanism underlying the protective effects of the exo-Gb3-FSL constructs. Coincubation of exo-Gb3-FSL, Stx1 and CHO-Lec2 A4G cells may trigger competition in Stx1 binding between cell-related receptors (Gb3Cer and P1 glycotope) and exosomes containing Gb3 trisaccharide.

Extracellular vesicles including exosomes play a key role in infectious disease pathogenesis, such as viral-, bacterial- or parasites-related diseases (e.g., HIV, SARS-CoV-2, *Mycobacterium tuberculosis*) [26, 27, 28]. Exosomes can modulate host immune responses by transporting pro- or anti-inflammatory molecules, thereby influencing the progression of infection. Furthermore, there are ongoing efforts to harness exosomes as potential therapeutic agents [29, 30], and for vaccine development [31, 32]. It has been demonstrated that exosomes are involved in Stx2 pathogenesis due to their ability to induce cell death through exosome-containing Stx2 [33, 34]. Future therapeutic strategies to combat Stx-related diseases in humans should be characterized by non-toxicity, non-immunogenicity, and target specificity. The host-derived exosomes may represent a promising approach for developing safe and personalized anti-Stx therapy. The introduction of exo-Gb3-FSL constructs offers a potential solution to the current limitations of Stx-related therapies; however, further investigations are necessary to evaluate their therapeutic efficacy in humans. Despite the need for additional research, exo-Gb3-FSL constructs appear to be promising new avenues for the development of innovative treatments for Stx-related diseases.

## Abbreviations

A4galt: α1,4-galactosyltransferase, Gb3/CD77 synthase
Exo-Gb3-FSL: exosomes containing incorporated FSL conjugate with Gb3 trisaccharide
Gb3: globotriaosylceramide, Gb3Cer, CD77, P^k^ antigen, Galα1→4Galβ1→4Glc-Cer
FSL: Functional-Spacer-Lipid
GP: glycoprotein
GSL: glycosphingolipid
HUS: hemolytic-uremic syndrome
STEC: Shiga toxin-producing *Escherichia coli*
Stx: Shiga toxin
Stx1B: Shiga toxin 1B subunit
Stx2B: Shiga toxin 2B subunit.

## Acknowledgments

The Fig. 4 was created with BioRender.com.

